# OpenComplex I: Predicting Three-Dimensional Atomic-Level Structures of Biomolecular Complexes

**DOI:** 10.1101/2025.03.25.643589

**Authors:** Jingcheng Yu, Zhaoming Chen, Zhaoqun Li, Mingliang Zeng, Wenjun Lin, He Huang, Qiwei Ye

**Affiliations:** Beijing Academy of Artificial Intelligence

## Abstract

OpenComplex^1^ is an open-source platform for comprehending and developing biomolecular models. Based on the architectures of AlphaFold2 and AlphaFold-Multimer, the current version of OpenComplex realizes end-to-end predictions of the biomolecular three-dimensional (3D) structures including protein, RNA, and protein-RNA complexes. OpenComplex makes it able to infer and train models of biomolecular complexes in a unified framework and supports encoders that use both MSAs and protein language models (pLM). Through extensive training on both structure and sequence databases such as the Protein Data Bank (PDB) and RNA families (Rfam) database, OpenComplex is able to generate models with equivalent or better accuracy for a wide range of biomolecules including monomers or multimers of protein and RNA as well as protein-RNA complexes. In summary, OpenComplex is a powerful tool for predicting accurate biomolecular structures in diverse application scenarios.

## 1 Introduction

Almost all biological processes in living organisms occur through specific interactions between biomolecular complexes, such interactions are therefore a crucial step in the investigation of biological systems and a starting point for drug design[Vangone et al., 2017]. One of the fundamental approaches to comprehending a biomolecule’s functions and the underlying chemical processes is to understand its 3D structure. The Critical Assessment of Structure Prediction (CASP) experiment has seen much progress in protein 3D structure prediction in recent years. For example, the representative method AlphaFold2 (AF2)[Jumper et al., 2021] showed accuracy competitive with experimental structures in most monomer instances, and end-to-end method AlphaFold-Multimer[Evans et al., 2022] achieved better performances compared with traditional protein docking methods in protein complex structure predictions. The reimplement of AlphaFold2 is made available by works like OpenFold[Ahdritz et al., 2022], Uni-Fold[Li et al., 2022], and ManyFold[Villegas-Morcillo et al., 2023]. Uni-Fold is currently the only open-source repository that supports the training of multimeric protein models. ManyFold proposed a proof-of-concept pLM-based model, pLMFold, which is trained from scratch to obtain reasonable results with reduced computational overheads in comparison to AlphaFold. These trainable, open-source codes are useful to understand how AlphaFold was organized and help advance methods for protein 3D structure predictions.

Similar to CASP, RNA-Puzzles provides a blind test for RNA 3D structure prediction tools[Cruz et al., 2012]. These evaluated tools can be classified into two groups: template-based structure assembly methods[Xu et al., 2014, Biesiada et al., 2016, Watkins et al., 2020] and physics-based folding simulation methods[Boniecki et al., 2016]. Recently, deep learning algorithms have also been used to design RNA structure prediction tools[Pearce et al., 2022, Shen et al., 2022]. For example, E2Efold-3D utilizes a framework similar to AF2 to predict RNA 3D structure and achieves an average sub-4Å root-mean-square deviation (RMSD) prediction accuracy on the RNA-Puzzles set[Shen et al., 2022]. However, the performance of current DL-based methods in CASP15 is significantly inferior to that of conventional physics- and template-based approaches because RNAs possess more flexible conformations than proteins[CASP, 2022].

Encouragingly, the implementation of OpenFold and E2Efold-3D demonstrates that it is possible to develop a unified framework for predicting the 3D structure of biomolecules such as proteins and RNA. Extending the RoseTTAFold[Baek et al., 2021] with a three-track layout, the newly released RoseTTAFoldNA[Baek et al., 2022] permits structural prediction of protein-nucleic acid complexes in addition to protein and nucleic acid monomers. Here, we proposed OpenComplex, an OpenFold-inherited AF2-style implementation of biomolecular tertiary structure prediction. This method allows for the prediction of protein and RNA monomers, as well as protein-RNA complexes, and produces comparable results to other prediction systems in the presence of homologous templates. We anticipate that OpenComplex will evolve into an AI system that is motivated by molecular structures and may be used in a variety of situations, such as structure design, in addition to being a tool for predicting the 3D structure of biomolecules in the future.

The rest of this report introduces the OpenComplex method and some experimental results. Since this is a long-term project, new results and codes will continue to be updated.

## 2 Methods

### Designing deep learning architectures for biomolecular complexes

We designed the deep learning architecture based on the layout of OpenFold[Ahdritz et al., 2022], which reimplements the AlphaFold2 architecture consisting of an MSA encoder and an atomic structural decoder (Figure 1). Given joined sequences as input, OpenComplex divides the sequence into multiple chains according to the biomolecular identifier and extracts the associated features such as MSAs and residue pairings for each chain type. Then a joined MSA and residue paring were generated by concatenating corresponding information. In addition, template structures and secondary structures were respectively retrieved for protein or RNA sequences and added as additional information to the residue pairing information. The co-evolution information represented by MSA and spatial constraints represented by parings were passed into the MSA encoder to extract the hidden representation of the input sequence, which is utilized in the structure module to direct the folding of the structure.

**Figure 1.**
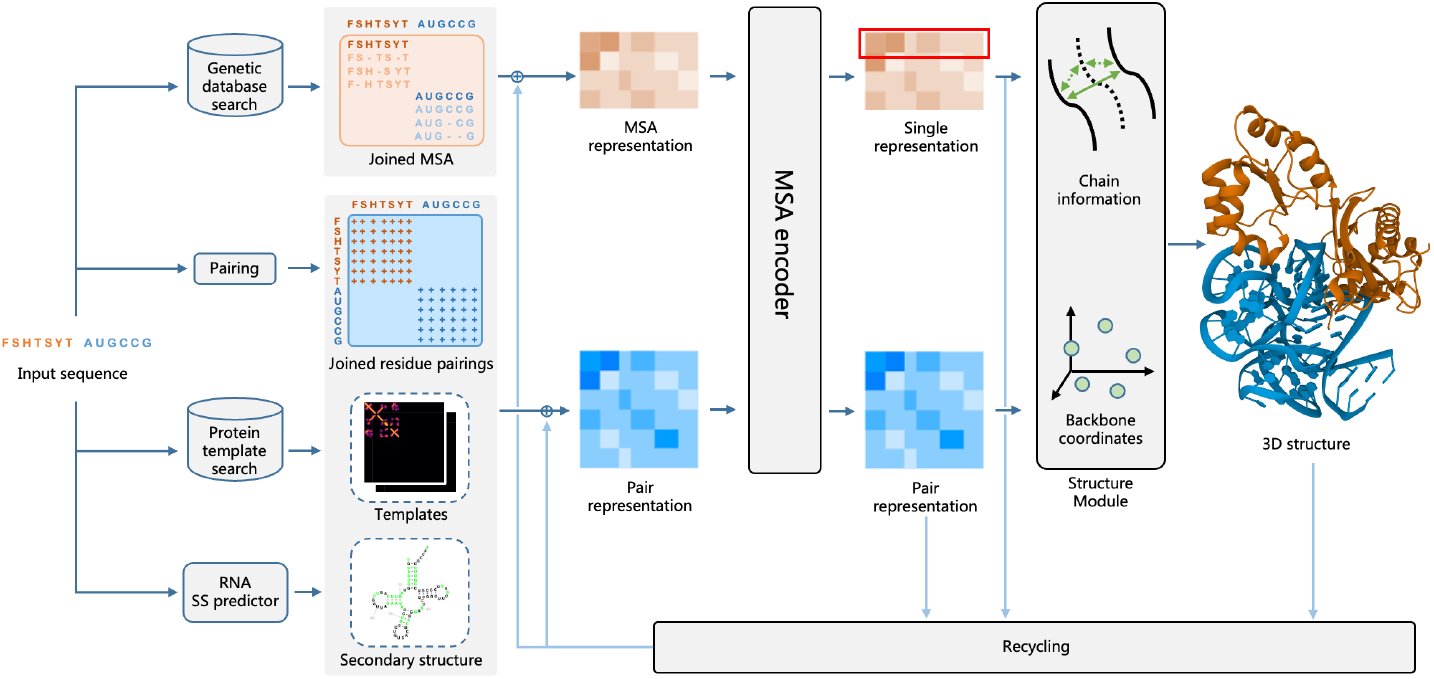
OpenComplex model architecture.

#### MSA encoder

The information on MSA representation and pair representation was updated using the MSA encoder, which is inherited from the Evoformer block proposed in AF2. To effectively leverage the positional information, the Rotary Position Embedding (RoPE)[Su et al., 2021] was utilized in the attention layer for both node representation and edge representation (Figure 2A).

**Figure 2.**
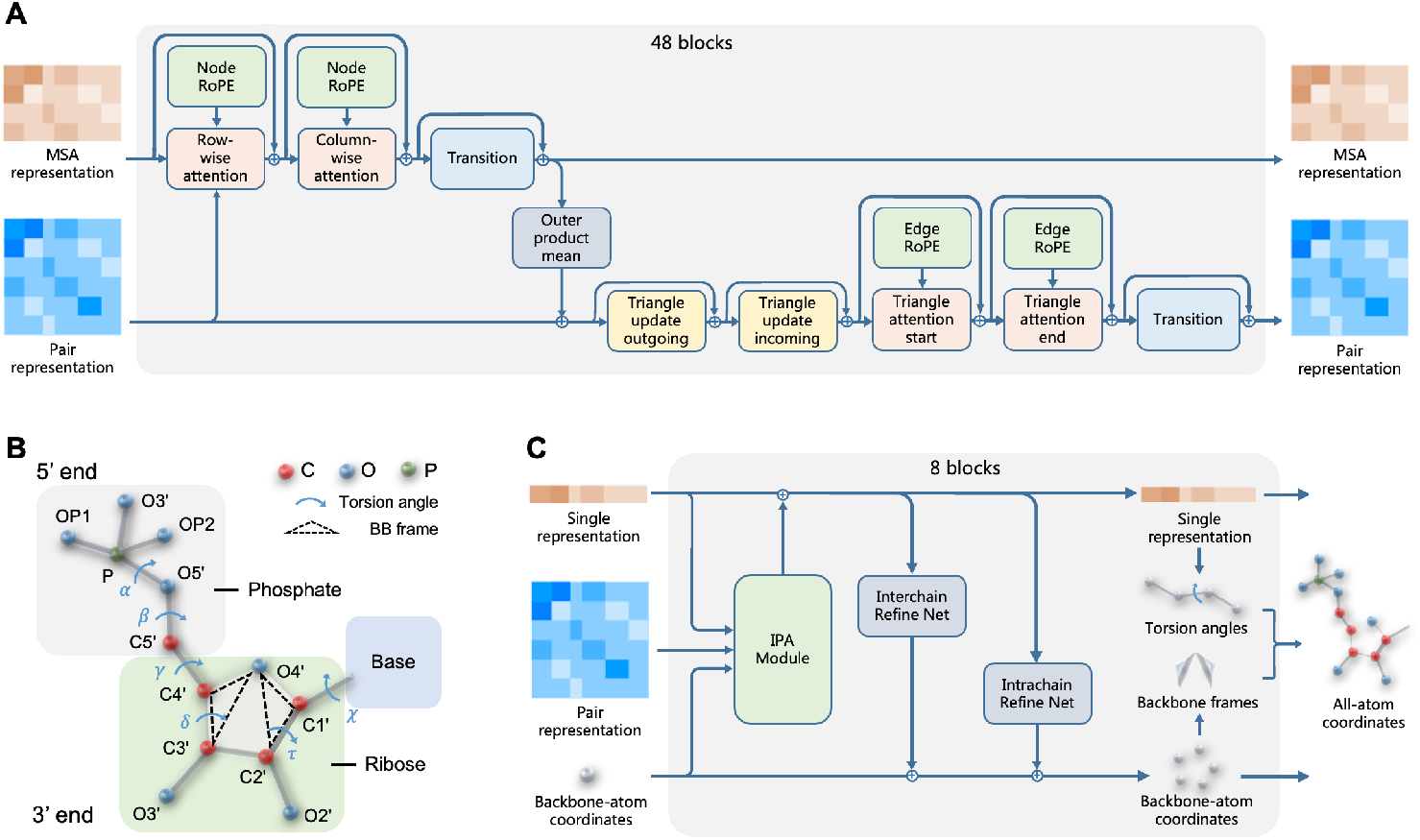
The schematic diagram of MSA encoder and structure module. A) MSA encoder block. RoPE, rotary position embedding. B) Schematic diagram of the atoms and torsion angles that make up a nucleotide. BB, backbone. C) Structure module block. Both intra-chain and inter-chain relative rotations and transformations were predicted and utilized for backbone-atom coordinates updates.

#### Designing molecule-specific frame representation in structure module

OpenComplex employs the same amino acid (AA) modeling method as AlphaFold2, which utilizes *N*, *Cα*, and *C* atoms to represent the AA backbone as a rigid frame and reconstructs side chain atoms using torsion angles. Unlike amino acids, the atoms in nucleotides are distributed in three subunit molecules: a ribose (five-carbon sugar), a phosphate group, and a nucleobase (Figure 2B). Since 5-membered rings are highly flexible, the 5 atoms in ribose are not coplanar, typically one or two atoms of the ring twist out of the plane, causing sugar pucker. The sugar puckers in DNA/RNA structures are predominately in either *C*3′-endo (A-DNA or RNA) or *C*2′-endo[Murray et al., 2003], making it difficult to represent the ribose with a single backbone frame. To solve this problem, we designated *O*4′, *C*4′, and *C*3′ as backbone frame 1 (bb1) to generate atoms along the phosphate direction, and designated *O*4′, *C*1′, *C*2′ as backbone frame 2 (bb2) to generate atoms along the nucleobase direction (Figure 2B). The stability of the bond angles *O*4′ − *C*4′ − *C*3′ and *O*4′ − *C*1′ − *C*2′ suggests that the ribose atoms we picked can be treated as rigid frames to represent the remaining atoms[Kowiel et al., 2020]. To get the all-atom representation of a nucleotide unit, seven torsion angles are needed besides the atom coordinates of ribose (Figure 2B). The incorporated seven torsions and related atoms are provided in Table 1.

**Table 1:** Rigid groups for constructing all atoms from given torsion angles. The *δ, γ*, and *α* torsion angles cooperate with bb1 to generate atoms along the phosphate direction, and the *τ*, *χ* torsion angles cooperate with bb2 to generate atoms along the base direction.

| type | bb1 | $\delta$ | $\gamma$ | $\beta$ | $\alpha_1$ | $\alpha_2$ | bb2 | $\tau$ | $\chi$ |
| --- | --- | --- | --- | --- | --- | --- | --- | --- | --- |
| A | O4', C4', C3', C5' | O3' | O5' | P | OP1 | OP2 | C1', C2', N9 | O2' | C4, N1, N3, N6, N7, C2, C5, C6, C8 |
| U | O4', C4', C3', C5' | O3' | O5' | P | OP1 | OP2 | C1', C2', N1 | O2' | C2, N3, C4, C5, C6, O2, O4 |
| G | O4', C4', C3', C5' | O3' | O5' | P | OP1 | OP2 | C1', C2', N9 | O2' | C4, N1, N2, N3, N7, C2, C5, C6, C8, O6 |
| C | O4', C4', C3', C5' | O3' | O5' | P | OP1 | OP2 | C1', C2', N1 | O2' | C2, N3, N4, C4, C5, C6, O2 |

The architecture of the structure module was modified to accommodate the processing of RNA units. Via the IPA module, the structure module in AF2 integrates the information in paired and single representations, leading the backbone frame update. Yet, because the two backbone frameworks of nucleotides share the *O*4′ atom in our design, direct prediction of the backbone frame will result in inconsistent *O*4′ coordinates. Thus for the nucleotide backbone, the structure module predicts the update for five atomic coordinates rather than for two rigid frames (Figure 2C).

### Preparing data for both monomeric and multi-chain molecule structures

We retrieved experimentally resolved structural data from the PDB database for protein/RNA monomers, protein/RNA multimers, and protein-RNA complexes. As of November 18, 2022, a total of 193,950 entries containing protein structure and 6463 entries containing RNA structure had been retrieved. This includes 182,652 entries of protein monomer structure, 1704 entries of RNA monomer data, and 4679 entries of protein-RNA complex data. After eliminating redundant sequences and sequences that are too long or too short, we divided the remaining data into the training and validation sets, in which the data released before May 1, 2020 was taken as the training set, and the data released after May 1, 2020 was taken as the validation set.

We generated features for protein monomers and multimers using pipelines consistent with AlphaFold2[Jumper et al., 2021] and AlphaFold-multimer[Evans et al., 2022], respectively. As for RNA sequences, the potential MSAs and secondary structures were fully explored together with relevant published experimental data. Briefly, MSA was generated using rMSA[Zhang et al., 2022] by querying genetic databases such as Rfam, RNAcentral, and nt databases, and the secondary structure was predicted from the MSA information using PETfold[Seemann et al., 2008]. For RNA multimers and protein-RNA complexes, the joined MSA and residue pairings were generated by simply concatenating features of the corresponding monomer sequence.

### Loss functions

The loss function of Opencomplex for protein monomers and multimers is inherited from AlphaFold2[Jumper et al., 2021] and AlphaFold-multimer[Evans et al., 2022]. The idea of the loss function for RNA structure is similar to that of protein structure, yet additional loss functions were introduced to constrain the intra-residue and inter-residue relationships. Specifically, intra-residue loss consists of *L*(*bb*_*dis*) and *L*(*bb*_*ang*) to constrain the bond length and bond angle of ribose, respectively, while inter-residue loss consists of *L*(*dis*_*geometry*) and *L*(*dis*_*torsion*) to constrain the distance and angle between adjacent residues, respectively. The introduction of additional intraresidue loss was necessary as we utilized two backbone frames to represent the ribose of the RNA residue, where *L*(*bb*_*dis*) is the average error between predicted and true distance for *C*2′ − *C*3′, *C*1′ − *C*4′, and *C*1′ − *O*4′, and *L*(*bb*_*ang*) is the average error between predicted and actual angles for *C*1′ − *C*2′ − *C*3′, *C*2′ − *C*3′ − *C*4′, and *O*4′ − *C*1′ − *N* 1*/N* 9. The inter-residue loss is similar to the energy function defined by DeepFoldRNA[Pearce et al., 2022], which measures the pairwise distance between adjacent *C*4′, *P*, and *N* 1*/N* 9 atoms, as well as inter-residue pseudo torsion angles. A log softmax activation is applied to the binned distance to calculate the error.

Gradients for protein and RNA structures are primarily derived from frame alignment point error (FAPE), however, RNAs and proteins differ slightly in how FAPE is calculated. As an RNA residue is represented by two backbone frames, those frames were concatenated along the chain dimension and then utilized to calculate the FAPE loss as defined in AlphaFold2.

The intra-residue torsion angles were measured by *L*(*supervised*_*chi*), which is the same as that defined in AlphaFold2.

Furthermore, we kept the *L*(*plddt*) loss introduced in proteins to measure the error between predicted lDDT (plDDT) and true lDDT. The parameter for binning is set to the same as those in proteins to discretize the lDDT value into 50 bins with an interval of 0.02, then a softmax crossentropy loss was calculated by comparing the binned lDDT value and binned plDDT value. The masked MSA error utilized in protein is removed for RNA as there are only four types of nucleotides.

### 3 Experiments

### Accurate predictions on monomeric RNA structures

We first evaluate the performance of OpenComplex in reconstructing RNA tertiary structures from primary sequences on the recently released CASP15 structures and RNA-Puzzles datasets. Notice that there are two tracks in CASP15 to submit predicted structures, including the Server track and the Human track. The server track demands the submission of predicted structures within 48h of the time when the sequence was released, while the human track provides one week of time for participants to design the structures. Herein, we evaluated all the submissions in both the server track and the human track.

#### CASP15

We chose the R1108 sequence from the chimpanzee CPEB3 ribozyme as a representative case due to its available ground truth structure (PDB ID: 7QR3). 26 groups participated in the submission for the human track, while only six groups submitted structures in the server track. By superimposing the predicted structures to the native structure, we calculated both root-meansquare deviation scores and TM-scores to evaluate model performance. Table 2 summarizes the statistical results for all the submitted and OpenComplex-generated structures. As presented, the structure predicted by OpenComplex achieves an RMSD of 3.55Å, outperforming five times better than the average RMSD of structures in both the human track and server track. The TM-score of the OpenComplex-predicted structure achieves 0.454. As a TM-score of 0.5 indicates the similarity of global folding among two structures, our predicted structure is far more similar to the native one than other structures. Since each group can submit at most five structures for a given sequence, we selected the best model of each group according to the RMSD to recalculate the average scores. Similar results were shown in Table 2. To further examine the results, we plotted the score distributions for all structures submitted in the human track. As is shown in Figure 3A, our predicted model achieves the best RMSD across 111 proposed structures, while the TM-score of our model outperforms most of the structures. For details, we superimposed our model and G489_4 to the native structure to examine the folding, respectively, as G489_4 achieves the best TM-score of 0.493. Figure 3B indicates that both structures are well-matched to the native structure in global folding. However, G489_4 did not capture the local structure of loop regions presented in red boxes, resulting in a poor RMSD value. Though possessing well RMSD, our model did not well understand the base parings of this structure (Figure 3C), especially in the P2 region.

**Table 2:** Quantitative comparison between OpenComplex predicted structure and submitted structures on CASP15-R1108.

| Method | RMSD (Å) | TM-score |
| --- | --- | --- |
| R1108_Human_all | 18.65 | 0.285 |
| R1108_Human_best | 16.04 | 0.302 |
| R1108_Server_all | 17.05 | 0.310 |
| R1108_Server_best | 14.52 | 0.302 |
| OpenComplex (ours) | 3.55 | 0.454 |

**Figure 3.**
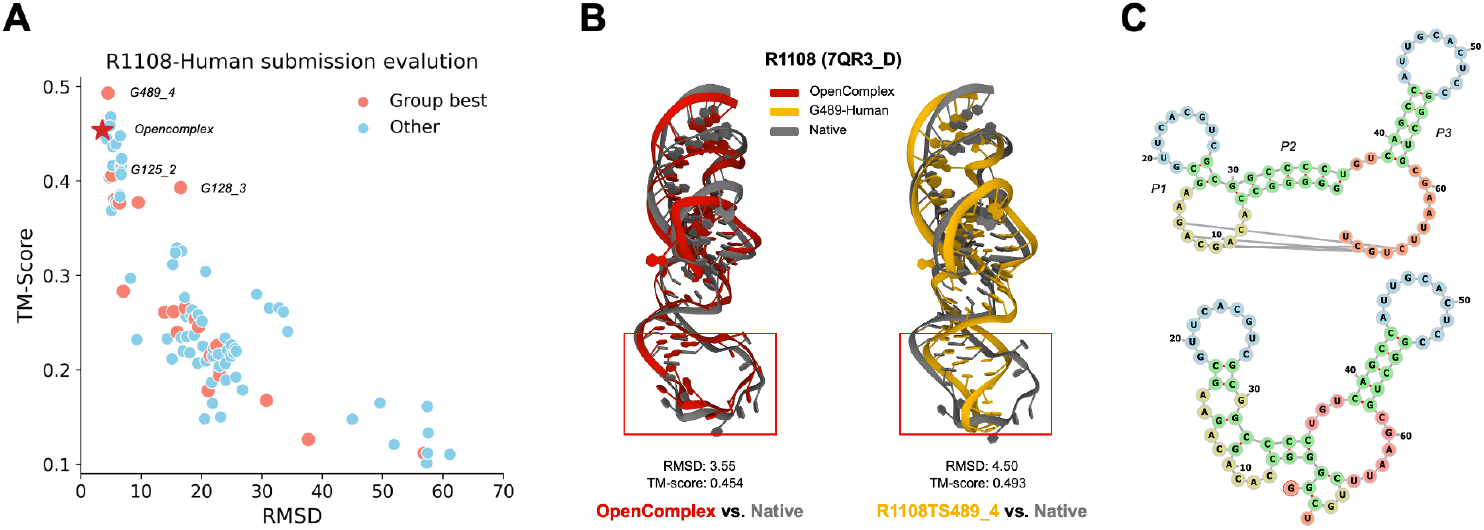
Comparison of performance of different methods on CASP15-R1108 structure. A) Distribution of evaluation metric scores for submitted structures in the human track. The x-axis stands for the RMSD score while the y-axis stands for the TM-score. The score of the structure predicted by OpenComplex was denoted with an asterisk. B) Visualization of the native structure and predicted structures of OpenComplex and Group-489. The native structure was shown in grey, OpenComplex predicted structure was shown in red, and Group-489 submitted structure was shown in yellow. C) Schematic diagrams of the secondary structure generated from the native structure (upper panel) and OpenComplex-predicted structure (lower panel).

#### RNA-Puzzles

Since the structures involved in RNA-Puzzles were eliminated from our training set, we could evaluate the performance of OpenComplex in predicting RNA monomers on the Puzzles data set. As a comparison, we sampled 500 structures for each sequence using FARFAR2 and selected representative structures based on ARES scores. DeepFoldRNA provides benchmarks for several RNA-Puzzles sequences, including refined and unrefined structures. We choose the unrefined structure as a comparison to our unrelaxed model. Furthermore, we selected the best submission for each RNA-Puzzles entry based on the RMSD score. As summarized in Table 3, OpenComplex achieves similar RMSD to DeepFoldRNA, and both deep learning approaches outperform FAR-FAR2 in model building, possibly due to the presence of homologous sequences.

**Table 3:** Quantitative comparison of structures predicted by OpenComplex with other methods on the RNA-Puzzles dataset. Missing values in DeepFoldRNA indicate that the structure is not in its provided benchmark.

| ID | Entry | Length | RNA-Puzzles<br>best submit | FARFAR2<br>+ ARES | DeepFoldRNA | OpenComplex<br>(ours) |
| --- | --- | --- | --- | --- | --- | --- |
| 4 | 3V7E_C | 126 | **3.257** | 14.27 |  | 5.8 |
| 5 | 4P9R_A | 189 | 9.15 | 20.77 | 3.38 | **2.77** |
| 7 | 4R4V_A | 185 | 20.37 | 14.7 | 3.53 | **2.02** |
| 8 | 4L81_A | 96 | 4.8 | 13.31 |  | **3.54** |
| 9 | 5KPY_A | 71 | 6.06 | 12.22 | 6.92 | **4.1** |
| 10 | 4LCK_B | 75 | **2.505** | 7.64 |  | 3.08 |
| 11 | 5LYS_A | 57 | 4.99 | 5.06 | 4.08 | **3.92** |
| 12 | 4QLM_A | 111 | 10.06 | 16.39 | **3.31** | 3.68 |
| 13 | 4XW7_A | 71 | 5.41 | 4.85 | **3.56** | 3.63 |
| 15 | 5DI4_A | 68 | 6.45 | 7.9 |  | **1.15** |
| 18 | 5TPY_A | 71 | 3.21 | 12.33 | **2.69** | 3.32 |
| 21 | 5NWQ_A | 41 | 3.65 | 9.27 | 2.28 | **1.69** |
| 23 | 6E8U_B | 37 | 8.13 | 7.62 | **3.31** | 4.24 |
| 25 | 6P2H_A | 69 | **2.55** | 8.99 | 3.2 | 2.67 |
| 29 | 6TB7_A | 52 | 4.3 | 6.01 |  | **1.04** |
| 33 | 7ELP_A | 46 | **3.4** | 3.5 |  | 3.46 |

### Accurate predictions on RNA-dimer structures

We also evaluate the performance of OpenComplex in reconstructing 3D structures of RNA multimers, from which we select two representative cases for analysis. The first case is 7TQV, which is a heterologous double-stranded RNA intermediate generated by coronaviruses during viral replication. The second case is 7TZS, which provides the crystal structure of the E. coli thiM riboswitch made up of two homologous sequences. We compared OpenComplex with RoseTTAFoldNA, which is the only available method for predicting structures of RNA multimer. As is shown in Figure 4A, both OpenComplex and RoseTTAFoldNA achieve a sub-3.5Å RMSD against the native structure of 7TQV, demonstrating the capability of the two methods in reconstructing the conformation of regular base pairings. However, the two methods exhibit different prediction patterns when predicting the structure of 7TZS, which is connected by GC pairings between two chains (Figure 4B). OpenComplex achieves a sub-2Å RMSD against the native structure, which benefits from accurate modeling of global conformation between chains as well as local secondary structure within chains (Figure 4B-C). However, RoseTTAFoldNA believes that more base pairing should be formed between the two chains, resulting in a totally different conformation against the native structure (Figure 4C).

**Figure 4.**
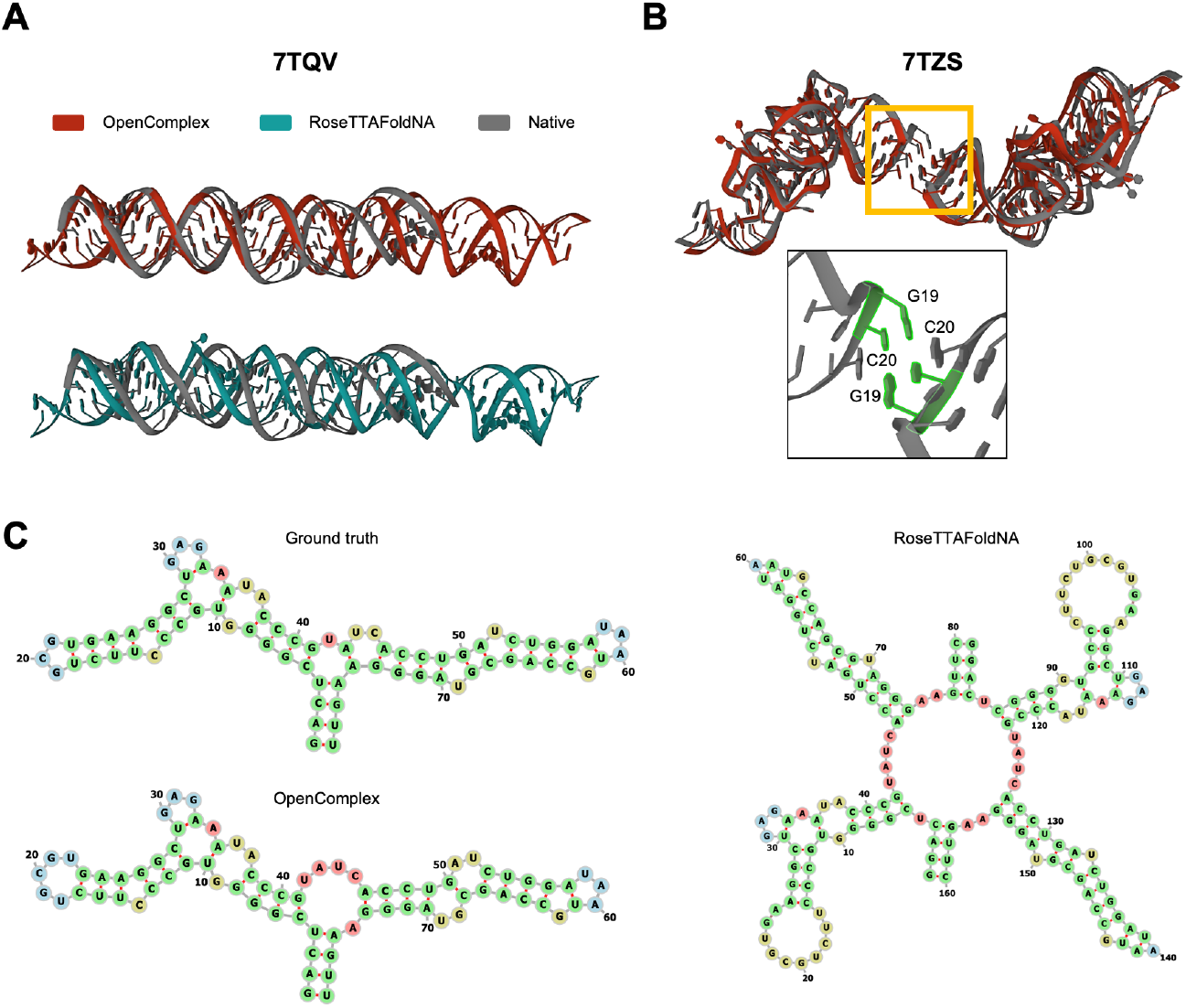
Detailed analysis of the results on RNA dimer structures. A) Visualization of the native structure for entry 7TQV and the predicted structures from OpenComplex and RoseTTAFoldNA. The native structure was shown in grey, OpenComplex predicted structure was shown in red, and the RoseTTAFoldNA predicted structure was shown in cyan. B) Visualization of the native structure for entry 7TZS and the predicted structure from OpenComplex. The native structure was shown in grey, Opencomplex predicted structure was shown in red. The zoomed area indicates the interaction interface formed by the GC base pairings. C) Schematic diagrams of the secondary structures retrieved from the 3D structures in the PDB format. The single-chain secondary structure was plotted for both ground truth and OpenComplex-predicted 3D structures, and the double-strand secondary structure was plotted for the RoseTTAFoldNA-predicted 3D structure.

### Accurate predictions on protein-RNA complexes

We then assess OpenComplex’s ability to predict the 3D structure of protein-RNA complexes. As shown in Figure 5A, the current version of OpenComplex achieves similar performance as RoseTTAFoldNA on a validation set of 36 structures, from which we selected 6XH3 and 7X34 for detailed comparison. 6XH3 is the co-crystal structure of HIV-1 TAR (trans-activation response RNA) in complex with TAR-binding proteins (TBPs) variant 6.3, whose arginine (R)-rich *β*2-*β*3 loop inserts into the major groove of TAR[Chavali et al., 2020] (Figure 5B). Although both Open-Complex and RoseTTAFoldNA accurately modeled the structure of TBP6.3, RoseTTAFoldNA mispredicted the interaction interface of TBP with TAR, resulting in a worse RMSD than OpenComplex (Figure 5B). 7X34 is the Cryo-EM structure of certain fragments of RNC-RAC complex in presence of Ssb from S. cerevisiae 2. The resolved structures include the C-terminal four-helix bundle domain (4HB) from Zuo1 and the ES12 (helix 44 of the 18S rRNA). As shown in Figure 5C, the positively charged lysine (K) is enriched in the *α*-helix formed by residues S335 to A350, which inserts into the negatively charged phosphate backbone of the major groove of ES12[Chen et al., 2022]. However, neither OpenComplex nor RoseTTAFoldNA do a good job of modeling the interface mediated by electrostatic interactions, even though there are more than 9000 homologous sequences for the RNA sequence with only 130 residues (Figure 5D). Since the RMSD for both RoseTTAFold and Opencomplex is greater than 35 Å, the incorporation of physics-based or force-field knowledge is of great significance in model training.

**Figure 5.**
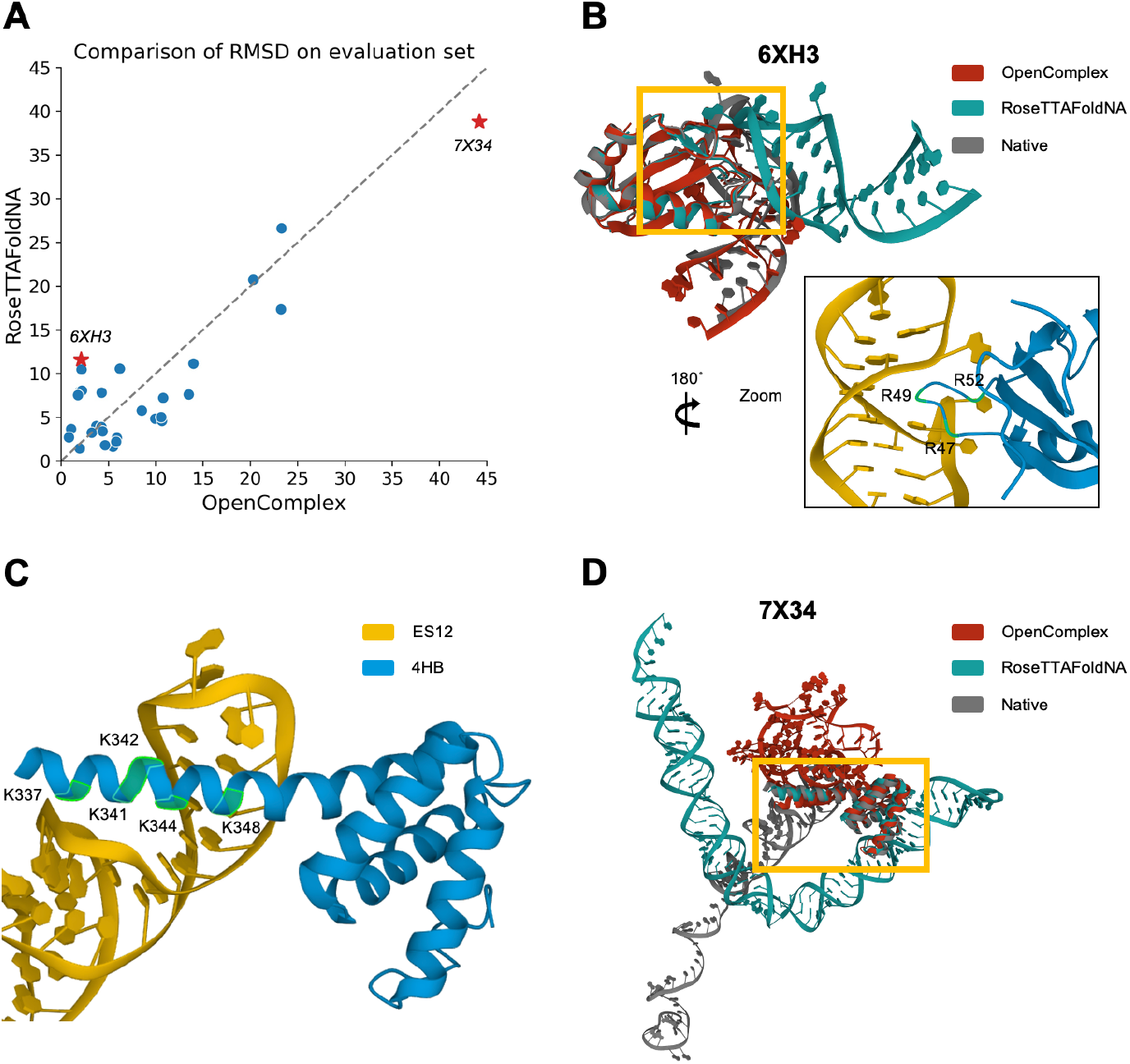
Detailed analysis of the results on protein-RNA complex structures. A) Comparison of the RMSD for OpenComplex and RoseTTAFoldNA on 36 evaluated protein-RNA complexes. The x-axis and y-axis stand for the RMSD score of OpenComplex and RoseTTAFoldNA, respectively. Asterisks mark two examples for detailed analysis. B) Visualization of the native structure for entry 6XH3 and the predicted structures from OpenComplex and RoseTTAFoldNA. The native structure was shown in grey, OpenComplex predicted structure was shown in red, and RoseTTAFoldNA predicted structure was shown in cyan. The zoomed area indicates the interaction interface formed by the R-rich *β*2-*β*3 loop in TBP6.3 and the major groove in TAR. C) Schematic diagrams of the interaction interface formed by the K-rich *α*-helix in 4HB and the major groove in ES12. The 4HB structure was shown in blue and the ES12 structure was shown in yellow. D) Visualization of the native structure for entry 7X34 and the predicted structures from OpenComplex and RoseTTAFoldNA. The native structure was shown in grey, OpenComplex predicted structure was shown in red, and RoseTTAFoldNA predicted structure was shown in cyan. The yellow frame marks the interaction interface.

## 4 Discussion and conclusion

The protein structure prediction tools represented by AlphaFold2 and AlphaFold-multimer have brought great changes to the development of structural biology. By transferring the corresponding ideas to nucleotide molecules, we realized the 3D structure prediction method of protein-RNA complexes and corresponding monomers and achieved a prediction accuracy similar to that of RoseTTAFoldNA. However, the current deep learning tools for the prediction of RNA 3D structure and protein-RNA complex 3D structure are not yet mature. Results for RNA tracks in CASP15 show that existing deep learning-based tools are far inferior to traditional physical or statistical force field-based tools when faced with scenarios lacking homologous sequences, such as artificially synthesized sequences. One of the solutions is to combine physics-based information. For example, Xiong et al. [2021] proposed a physics-based energy function that captures the full orientation dependence of base-base, base-oxygen, and oxygen-oxygen interactions of RNA 3D structures. The proposed method, named BRiQ, performed well in structure refinement and achieved the best average performance in the CASP15 competition.

Furthermore, the lack of labeled data and a greater degree of flexibility make it harder for RNA and protein-RNA complex structure prediction. We believe there should be a lower-level mechanism that can be modeled by molecule dynamics, with which to compensate for the gap between protein, RNA, and DNA modeling. Ramaswamy et al. [2021] uses force-field-based loss functions in the deep learning method to predict the possible state transition path in protein conformation space. ProtMD[Wu et al., 2022] shows a self-supervised pre-train model based on the MD trajectories that leads to SOTA downstream binding affinity prediction and ligand efficacy prediction tasks. These recent researches show molecule dynamics modeling could bring new chances for predicting protein conformation switches, biomolecular interactions, etc., which will further lead to better functionality prediction, and *de novo* designing.

## Footnotes

1 TCode is available at https://github.com/baaihealth/OpenComplex. OpenComplex is a work-inprogress project. The content of this technical report will be updated continuously

## Notes

https://github.com/baaihealth/OpenComplex

### Competing Interest Statement

The authors have declared no competing interest.

